# Cellular aspects of gonadal atrophy in P-M hybrid dysgenesis

**DOI:** 10.1101/041178

**Authors:** Natalia V. Dorogova, Elena Us. Bolobolova, Lyudmila P. Zakharenko

## Abstract

Gonadal atrophy is the most typical and dramatic manifestation of intraspecific hybrid dysgenesis syndrome leading to sterility of *Drosophila melanogaster* dysgenic progeny. The P-M system of hybrid dysgenesis is primarily associated with germ cell degeneration during the early stages of *Drosophila* development at elevated temperatures. In the present study, we have defined the phase of germ cell death as beginning at the end of embryogenesis immediately following gonad formation. The early stages of germ cell formation in dysgenic flies showed sensitivity to developmental temperature increases at any stage of the *Drosophila* life cycle including the imago. Analysis of germ cell reactions to hybrid dysgenesis induction revealed significant changes in subcellular structure, especially mitochondria, prior to germ cell breakdown. The mitochondrial pathology can be the reason for the activation of cell death pathways in dysgenic germ cells and lead to gonadal atrophy.

**Summary:** We investigated the hybrid dysgenesis in the context of germ cell development and observed the dysgenic effect is associated with dramatic pathological changes in structure of germ cell mitochondria and lead to massive and quick cell death. The temperature-dependent screening of germ cells developmental pattern in dysgenic background showed that these cells are susceptible to the hybrid dysgenesis at any drosophila life-cycle stage, including in the imago.

## Introduction

Crosses between some pairs of *Drosophila melanogaster* strains result in the sterility of hybrid offspring in one cross direction, which is most clearly manifested in females. It is believed that such events are the result of interaction of the **p**aternal (P) genome with the **m**aternal (M) cytoplasm. Therefore, this type of intraspecific gonadal dysgenesis (GD) is referred to as PM GD or PM hybrid dysgenesis (PM HD) (Sved, 1976; Kidwell et al., 1977). Such PM GD is susceptible to developmental temperature and is strongly displayed when dysgenic flies are raised at restrictive temperatures of 25– 29 °C. Lowering the temperature to 20 °C usually completely prevents PM GD (Engels, 1979; Kidwell and Novy, 1979).

Dysgenic gonads contaiSn an extremely reduced number of germ cells (GCs), however, a normal somatic background is maintained (Kidwell and Novy, 1979; Engels and Preston, 1979; Bhat and Schedl, 1997). The mechanism leading to GC absence is unknown; nevertheless, different causes are assumed including loss of specification, abnormal cell division, cell cycle arrest due to multiple DNA breaks, and others (Engels, 1996; Bhat and Schedl, 1997; Majumda and Rio, 2015). However, to understand the underlying mechanisms of this phenomenon it is necessary to determine what cellular event or process is the primary target and which developmental stage is the most sensitive. Therefore, to understand why the germ cells do not survive in the dysgenic background we monitored their development cytologically during the fly life cycle under various temperature conditions.

We focused on GC fates in the PM HD system induced by crossing females from the M-strain (M cytotype) with males from the P-strain (P-factor). In this study, we show that GCs of dysgenic hybrids participate in embryonic gonad formation; however, most subsequently die while the somatic cells of the gonad survive. Nevertheless, the phase of GC death is not exclusively embryonically determined and can occur during any stage of the *Drosophila* life cycle by altering the temperature conditions. In particular, we show that if the dysgenic flies are raised at 20°C and then moved to 29°C, their germ cells begin to lyse quickly en masse and then (during 3–5 days) die. However, pathological changes in the cytoplasm were observed most notably with the appearance of a large number of damaged mitochondria prior to germ cell death. We propose that cell death pathways are activated by mitochondrial dysfunction, and that the mitochondrial defects may result from nuclear-cytoplasmic conflict in the P-M system of HD.

## Materials and methods

### *Drosophila* culture and strains

We used *D. melanogaster* strains: *Canton S* obtained from I. Zakharov, Novosibirsk, Russia (maintained since 1968); *Harwich-w* obtained in 1996 from L. Kaidanov, Saint-Petersburg, Russia, and saved in the laboratory of Prof. I. Zakharov from Novosibirsk.

Control and dysgenic crosses were set up at the permissive temperature of 20 °C and at the restrictive temperatures of 25 and 29 °C.

### Electron and fluorescence microscopy

Experimental procedures for electron and fluorescence microscopy were performed as described previously (Pertceva et al., 2010).

Antibodies used were monoclonal anti-VASA (Santa Cruz Biotechnology; 1:300), anti-Fasciclin III (7G10, DSHB Hybridoma Product; 1:80), polyclonal guinea pig anti-Traffic Jam [kindly provided by Prof. D. Godt (Li et al., 2003), 1:400], Alexa-488-conjugated anti-guinea pig and anti-mouse IgG and Alexa-568-conjugated anti-rabbit IgG (Molecular Probes/*Invitrogen;* 1:400). We used *ProLong Gold* anti-fade reagent with DAPI staining (Molecular Probes/*Invitrogen)*.

For embryonic antibody staining, embryos were removed from agar plates with a paintbrush and dechorionated in 50% bleach for 5 min, followed by a cold water rinse to remove the bleach. Embryos were then fixed in 3.7% formaldehyde and heptane, devitellinised with methanol/heptane and stained with antibodies using standard protocols (Patel, 1994). Lysosome staining was carried out in 1 μM LysoTracker red DND-99 (Invitrogen, Molecular Probes, Basel, Switzerland) according to a previously described procedure (Dorogova et al., 2014). Embryonic germ cell development was analyzed in nearly 500 embryos, and later developmental stages were analyzed in no less than 100 flies per experiment.

Images were obtained using an AxioImager Z1 microscope with ApoTome attachment (Zeiss), AxioCam MR and AxioVision software (Zeiss, Germany).

## Results and Discussion

P-M HD results in full female sterility and a half-sterility of males raised at 29 °C. Dysgenic flies raised at this temperature had reduced gonads with an absence or significantly reduced number of germline cells. However, infertility can be prevented partially or completely when the developmental temperature is lowered to 25–20°C (Engels and Preston, 1979, Kidwell and Novy, 1979). We used the temperature effect on the fate of GCs to identify what stage (or event) of their development is targeted in dysgenesis.

To generate offspring with dysgenesis syndrome we crossed *CantonS* females (M strain) to *Harwich* males (P strain). The first control was the *CantonS* strain because these females are a source of germ plasm, and the second control were the offspring from a reciprocal cross because they are genetically identical to dysgenic F1. However, the data presented result from only one control because their comparison did not show significant differences.

We have conducted a detailed cytological analysis of the cell organization and morphology in dysgenic gonads subjected to alterations in the incubation temperature from restrictive to permissive (temperature shift) at different stages of the *Drosophila* life cycle. Cells were visualized with antibodies against the VASA protein to mark the germline and antibodies against the Traffic Jam and Fasciclin III proteins to mark soma.

Initially, to clarify how the dysgenic gonadal phenotype is generated by our genetic material we repeated earlier temperature shift experiments (Engels and Preston, 1979; Bhat and Schedl, 1997). The result we observed is generally consistent with the previous data (supplementary material Fig. S1). Based on this information, we have concluded that GC viability is not determined during specific events of embryogenesis (specification, proliferation, or migration), but immediately as the gonads form (supplementary material Fig. S1A, B). Mitigation of dysgenic conditions (25 °C) allows some GCs to be rescued in embryogenesis but they remain susceptible beyond this stage ( supplementary material Fig. S1C-H).

### Hybrid dysgenesis affects early germ cells, regardless of the ontogenetic stages

Since GCs are eliminated in already formed gonads, we assumed that this effect can be promoted at any stage of the fly life cycle. For providing HD conditions, dysgenic offspring of different developmental stages (larva, pupa or imago) were placed from 20 to 29 °C. Maintenance of dysgenic offspring at 20 °C beforehand avoids the effects of hybrid dysgenesis and provides normal gonad development.

Altering the developmental temperature from 20 to 29 °C for larvae and pupae induced germ cell degradation. GC reduction was most apparent on the 3rd day of incubation at the restrictive temperature (Fig. 1 A–D). Partial loss of GCs were observed in all gonads, however, only 3% of the females and 1% of the males demonstrated complete atrophy of germline tissue (n = 100).

**Figure 1.**
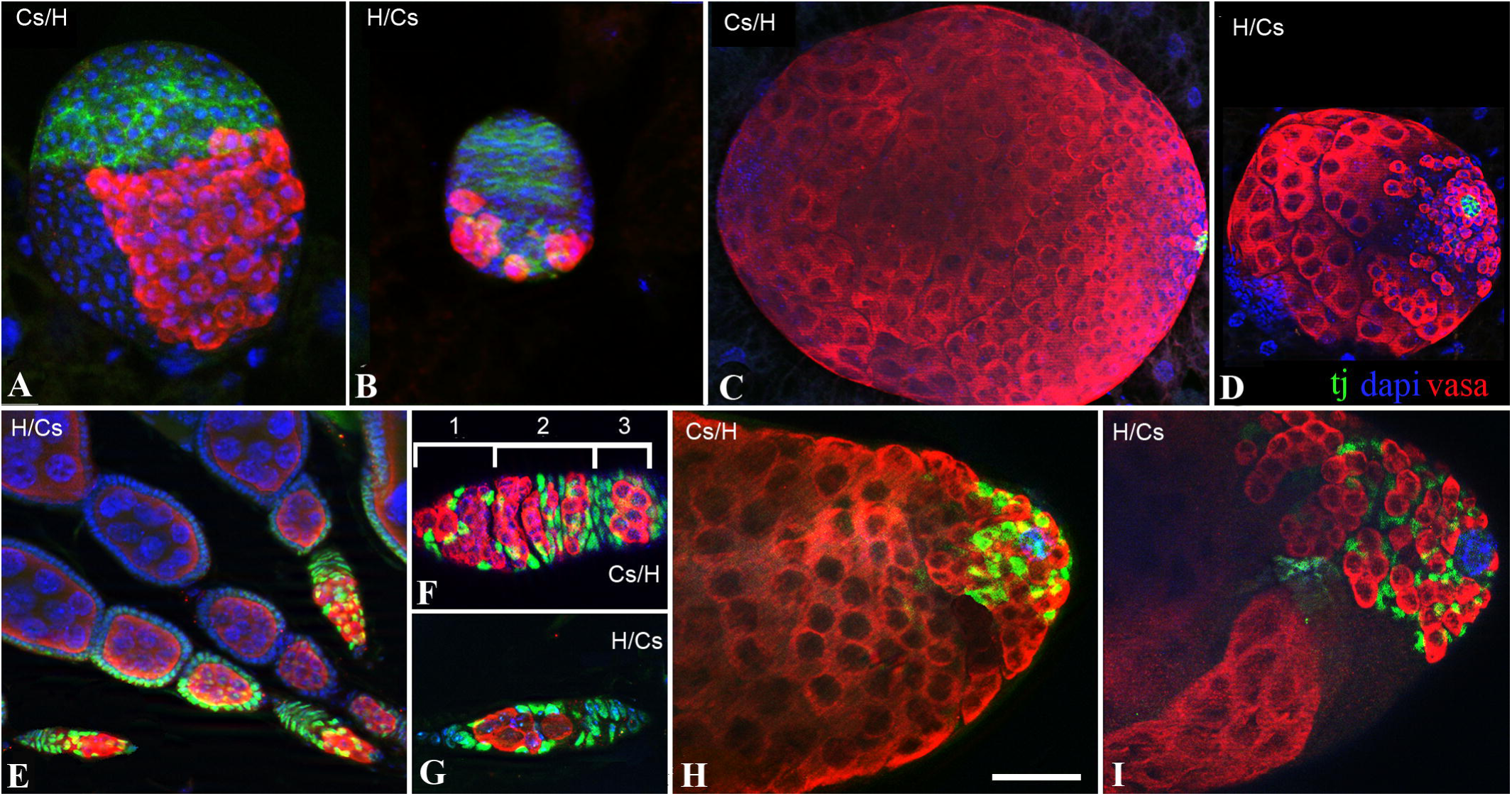
Dysgenesis-induced GC degeneration under developmental temperature variation from 20 to 29 °C. **A–D.** Larval gonads after incubation at 29 °C for 3 days. **A.** Control larval ovaries have a normal GC pattern **B**. Dysgenic ovaries are much smaller and contain very few GCs. **C,D.** Larval dysgenic testes (D) are reduced in both size and number of GCs compared to the control (C). **E-I.** Imago gonad after incubation of newly hatched flies at 29 °C for 3 days. **E.** Dysgenic imago ovarioles contain reduced germaria but normal egg chambers at later stages. **F,G**. Comparison of germaria in control (C) and dysgenic (D) ovaries. Dysgenic germaria contain significantly fewer germ cells than in the control and they are not differentiated in the regions of 1-2A-2B-3. **H.** In the control testis, the anterior end is completely filled with early GCs. **I.** In dysgenic testes, there are clearly identified VASA-negative areas lacking GCs. Germ cells are visualized by anti-Vasa (red), somatic cells – anti-FasIII (green), DNA - DAPI (blue). Scale bar: 5 μm (A, B); 10 μm (C-D); 20 μm (F-I).

In 1-day-old flies reared at 20 °C followed by the restrictive temperature for 3 days, the dysgenic gonads showed general germ cell depletion in the germarium of the ovaries and apical end of the testes where early-stage cysts are located (Fig. 1 E–I). Most later-stage GCs were resistant to the HD effect, and they developed normally to produce eggs and sperm. Females maintained at 29 °C for one week had significantly reduced reproductive systems: completely atrophied germaria and also significantly depleted vitellaria resulting from the inability of the germaria to produce new egg chambers. Males that developed in the same conditions also demonstrated a depletion of GC cysts especially in the anterior region of the testes. All of the analyzed dysgenic gonads (n∼200) showed significant and obvious early-stage GC loss.

Thus, the temperature increase is a trigger of hybrid dysgenesis phenotype regardless of the stage of the *Drosophila* life cycle, and germ cells in the early stages of their development are the most sensitive. Embryonic gonads consist exclusively of early-stage GCs and consequently undergo rapid atrophy.

### Hybrid dysgenesis induces massive germ cell lysis

The temperature-dependent screening of gonadal phenotypes showed that HD can cause significant early GC loss at any stage of fly development. To determine whether GCs die en masse and when the process begins, lysosomes were stained with LysoTracker. Regardless of the cell death pathway, there is a subsequent increase in lysosomal activity that can be detected by LysoTracker (LT) dye (Fogel et al., 2012).

Dysgenic 1-day-old flies and 1^st^ stage larvae were incubated at 20 °C and then transferred to 29 °C. The lysosomal activity in their gonads was monitored after 3 hours, 1 day, and 3 days incubation at the restrictive temperature. In all variants, the LysoTracker-labelled area was wider in dysgenic progeny than in controls (n = 100). In more detail, we have presented here an analysis of this effect in imago ovaries, because in this case, the pattern of lysosomal activity can be most clearly mapped relative to the inner structure of the ovary and easily quantified.

Lysosomal staining was observed in the ovaries of flies grown at 20 °C, as in dysgenic and control crosses, but did not exceed 20% of the germline cysts. Raising the temperature for 3 hours resulted in an increase in the number of LT-stained areas in control flies around 15%, in region 2a / 2b of the germarium and in mid-oogenesis. However, in dysgenic ovaries, an increase in the LT signal was detected in 95% of the ovarioles (n = 500), occupying not only 2a/2b but also other regions of the germaria (Fig. 2 A, B). In addition, lysis appeared in control ovaries locally and affected individual cysts. In dysgenic ovaries, only single cysts escaped death and could advance to the next stage of development. Most GCs did not migrate out of the 2A region of the germarium. After a 3-day incubation at 29 °C, the germaria were considerably reduced due to the loss of not only individual GCs but also regions 2B and C (Fig. 2 C, D). Egg chambers of later stages, including vitellogenesis, exhibited only slight degradation in comparison to the control.

**Figure 2.**
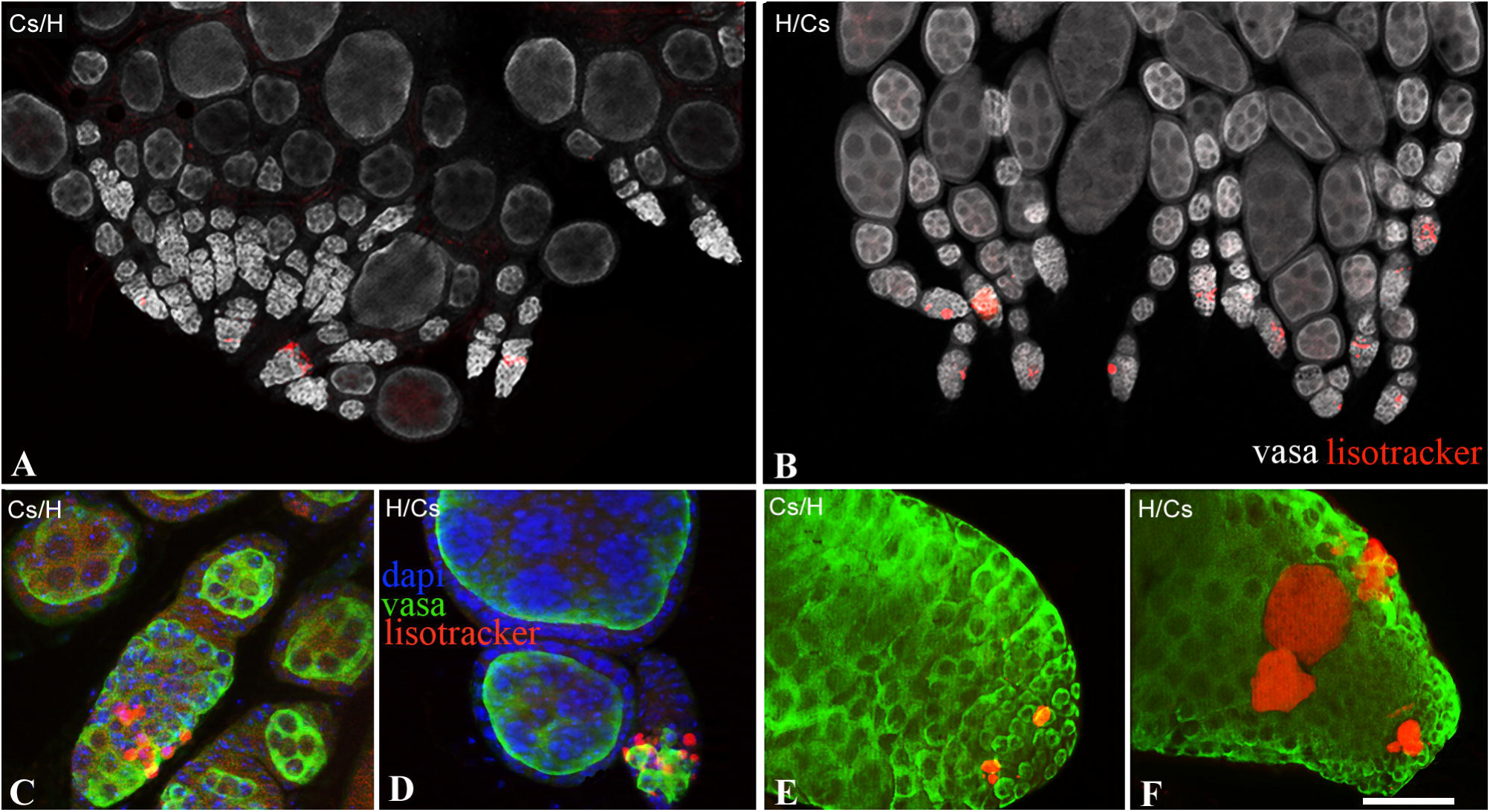
Lysis of germ cells, caused by hybrid dysgenesis. **A,B.** The ovaries of 1-day-old females after a 3-hour incubation at 29 °C. LysoTracker staining is observed in almost all germaria in dysgenic ovaries (B) in contrast to the control (A).**C-F.** GC lysis after a 3-day incubation at 29 °C. **C.** In the control, LysoTracker stains only a local area of the germarium, while retaining their normal structure and GC status **D.** In dysgenic ovaries, germarial regions 2B-3 are absent, and the remaining GCs are covered by lysis. **E,F.** Testes of control (E) and dysgenic (F) males after a 3-day incubation at 29 °C. There is massive lysis in dysgenic testes. Germ cells are visualised by anti-Vasa (green), DNA - DAPI (blue), lysosomes – LysoTracker (red). Scale bars: 50 μm (A,B); 15 μm (C–D); 10 μm (E-F).

Enhanced lysosomal activity was also observed in testis, mainly in the areas occupied by early-stage cells (Fig. 2 E, F).

### Hybrid dysgenesis significantly affects the ultrastructure of germ cells and causes extensive cell death

We used electron microscopic analysis to characterize the cellular pathology associated with the massive lysis under conditions inducing HD. The gonads of adult dysgenic flies that had developed at 20 °C had an ultrastructure similar to that of the control. Signs of the ultrastructural pathological changes were first visible after 3 hours of incubation of 1-day-old flies at 29 °C.

After a 3-hour incubation in females, in addition to increased number of lysosomes, we observed defects in mitochondrial morphology. In prefollicular GCs, mitochondria are readily detectable in Bolbiani bodies where they pass through the fusome. Abnormal mitochondria of dysgenic ovaries were less structured, had areas with electron light matrix and partially lacked cysts (Fig. 3 A, B, A’, B’).

**Figure 3.**
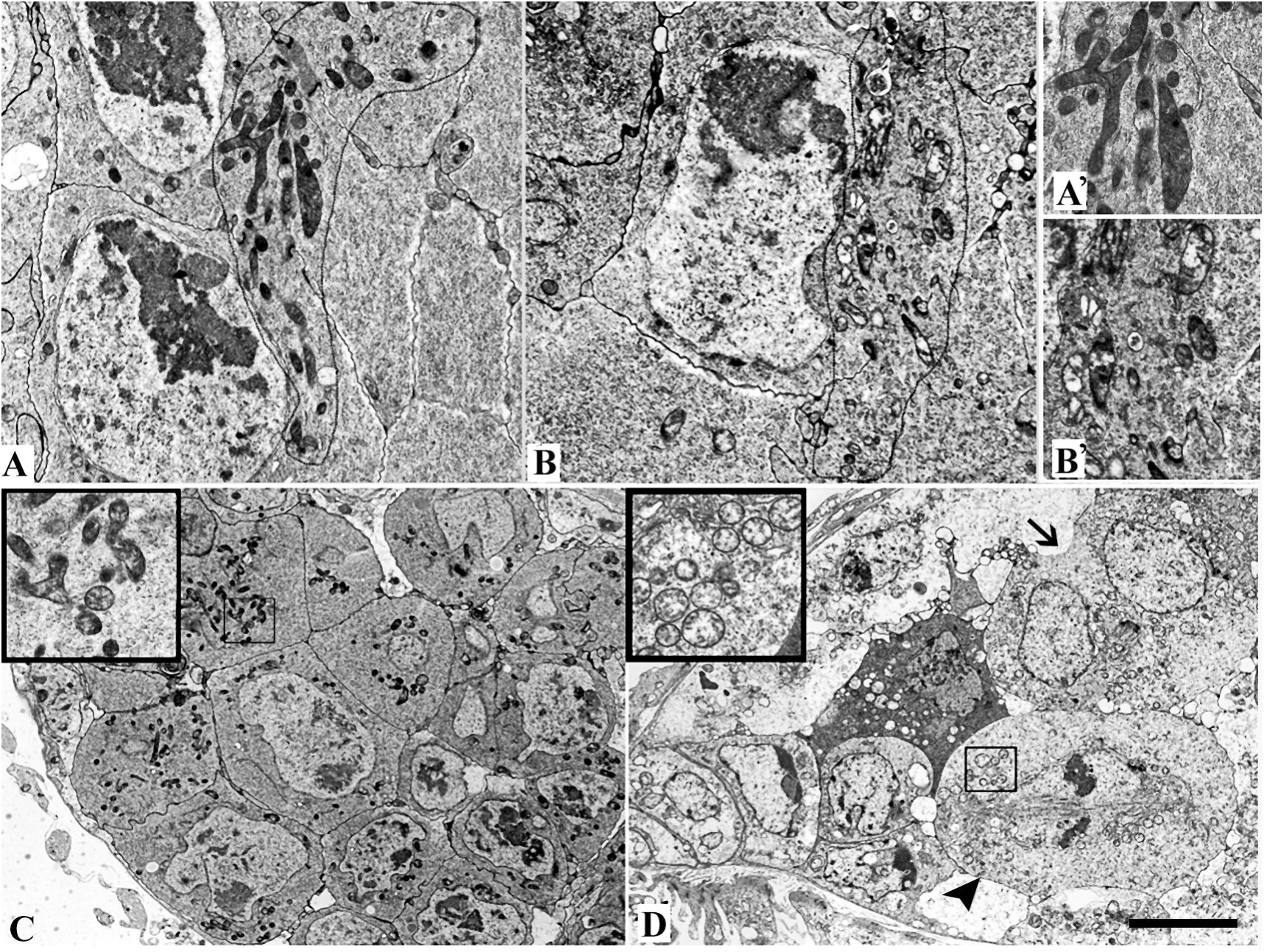
Anomalies in fly imago gonadal ultrastructure caused by hybrid dysgenesis. **A,B**. Internal GC structure and mitochondrial morphology in the 2A region of the germarium after a 3-hour incubation at 29 °C. Compared with the control (A), the GCs of dysgenic germaria are filled with defective mitochondria (B). **A’,B’.** Dysgenic mitochondria (A’) are significantly smaller than control mitochondria (B’), have areas with electron light matrix and are partially devoid of cristae. **C,D**. The ultrastructure of the 1-2A germarium region in the control (C) and dysgenic (D) females after a 3-day incubation at the restrictive temperature. Dysgenic germaria have signs of intracellular structural degradation and cell death. However, some cells continue to divide despite the pathological changes. Figure 4D demonstrate cells in metaphase (arrowhead) and anaphase (arrow). They form a normal cell division spindle and cleavage furrow, however the cytoplasm appears depleted and poorly granulated and mitochondria (highlighted in the square frame) are swollen and transparent. Scale bars: 0.3 μm (A–B), 0.2 μm (A’,B’), 1μm (C–D).

Increasing the incubation time at 29 °C for 3 days was associated with a significant expansion of the lytic areas and a reduction in the number of viable GCs (Figs. 3 C, D). This is consistent with the light microscopic data. However, viable and dividing germ cells also showed an enhancement of the cytoplasmic defects. There were poorly granulated cytoplasm, swollen mitochondria with transparent matrix and depleted cysts. However, these cells were able to form structures providing cell division, such as the division spindle and cleavage furrow (Fig. 3 D).

Dysgenesis induced anomalies in spermatogenesis phenotypically manifested as well in the oogenesis but to a lesser extent. We observed similar abnormalities in mitochondrial ultrastructure after a 3-hour incubation of males at 29 °C, but compared with oogenesis they were not widespread. After a 3-day incubation at 29 °C, the mass death of cysts in the testis was evident but it was not as large-scale as in the ovaries (not shown).

### Why are germ cells dying?

Programmed cell death is essential for normal germline development and plays a role in removing impaired or excess cells (Denton et al., 2013). In a typical *Drosophila* gonad, entire germline cysts undergo execution rather than single cells. In wild-type ovaries, GC death sporadically occurs after passing certain checkpoints within the germarium (region 2) and during mid-stages of oogenesis (stages 7–9) (McCall, 2004). During normal spermatogenesis, some of the spermatogonial cysts undergo spontaneous cell death (Yacobi-Sharon et al., 2013). In addition, checkpoint control provides selective removal of cells with signs of unrepaired defects; however, in the case of HD, a massive loss of early germline cysts was detected in our analyses. These phenotypes are similar to those observed after the overexpression of the effector caspase Dcp-1 and downstream mediator Dmp53, which are important in cell death pathways (Barth et al., 2011). Therefore, it is possible that the uncontrolled GC death in dysgenic backgrounds is not associated with the checkpoint system.

The mechanism promoting GC degradation in HD is not clear. However, the sudden appearance of numerous defective mitochondria would lead to problems with cell metabolism. Visually, the mitochondria are very first subcellular structures that react to the effects of HD, and their internal structure is considerably degraded during the several hours of incubation at the elevated temperature. It is known that mitochondria accumulate some key regulators of cell death such as *cytochrome c* and other pro-apoptotic proteins as well as dangerous reactive oxygen species (Krieser and White, 2009; Abdelwahid et al., 2011). Mitochondrial defects can cause disordering of the selective permeability of the outer membrane and the release of these factors into the cytosol triggering cell death.

## Conclusion

We investigated the HD in the context of GC development and observed that these cells become susceptible to the dysgenic effect only after embryonic gonad formation. However, manipulating the incubation temperature can induced the HD symptoms at any life-cycle stage of the dysgenic progeny, including in the imago. The gonads of adult dysgenic flies that had developed at 20 °C displayed a normal morphology and were filled completely with GCs. However, the displacement of these flies to the restrictive 29 °C led to mitochondrial defects in germ cells and promoted their lysis culminating in death. Quick and mass GC death creates difficulties in the study of molecular mechanisms of HD and experimental evidence of *P*-elements activity in induction dysgenic syndrome. Our research has shown that the GC lifetime can be controlled by means of temperature regulation. There is a lag-period between the beginning of HD induction and the onset of GC death. This time gap provides an opportunity to isolate the germline for molecular and cytological studies of *P*-element activity or other potential agents of HD.

## Author contributions

N.V.D. designed and conducted experiments, analyzed data, wrote and prepared the manuscript, E.Us.B and L.P.Z. conducted experiments and analyzed data.

